# Genome assembly for *Amanita muscaria* var. *muscaria*

**DOI:** 10.64898/2026.11.17.767403

**Authors:** Lydia Turley, Matt Larcombe, David Bryant, Augustine Chen, Janice Lord, Andy Nilsen, David Orlovich, Ludovic Dutoit

## Abstract

*Amanita muscaria* is a charismatic and widely distributed mushroom-forming fungus. *Amanita muscaria* comprises a species complex, and genomic resources are unevenly distributed, which may bias genetic and genomic study of the species complex. This study produces a high quality hybrid genome assembly of a specimen from New Zealand. Phylogenetic analysis shows that this belongs to *Amanita muscaria* var. *muscaria*, making this the first genome assembly for this group.

## 1. Introduction

The global success of invasive trees such as *Pinus* have often been enabled by cointroduced ectomycorrhizal fungi, which may themselves become invasive species (Hoeksema et al., 2020; Policelli & Nuñez, 2025; Vargas, Gonçalves, Franco-Molano, Restrepo, & Pringle, 2019). Yet in contrast to their hosts, exotic ectomycorrhizal fungi have received less attention and their impacts as invasive species are less well known (Díez, 2005; Policelli & Nuñez, 2025). This is at least partly due to the detectability of these mostly subterranean fungi (van Galen et al., 2025). Their cryptic life style makes understanding simple ecology and population demography challenging, and genomic approaches offer the best strategy to study these systems (Vincenot & Selosse, 2017). The development of high quality (highly contiguous) reference genomes for ectomyc-orrhizal fungi will facilitate the development of genomic tools to shed light on these invasion processes as well as ecological and evolutionary aspects of ectomycorrhizal relationships in general (Hoeksema et al., 2020).

*Amanita muscaria* is probably the most well-known mushroom-forming fungus globally, and is an important ectomycorrhizal species that has been widely co-introduced with exotic trees around the world. This charismatic species typically produces large red sporocarps with white spots that are remnants of the universal veil (Carboué & Lopez, 2021). *Amanita muscaria* is particularly associated with pine (*Pinus*), birch (*Betula*) and spruce (*Picea*), and is known to form associations with a number of other plant species (Hall, Stephenson, Buchanan, Yun, & Cole, 2003). It is native to Northern hemisphere temperate and boreal forests (Geml, Laursen, O’neill, Nusbaum, & Taylor, 2006) and has been recorded invading native forests from exotic plantations across the southern hemisphere, including New Zealand (Johnston, Buchanan, Leathwick, & Mortimer, 1998), Australia (Fuhrer & Robinson, 1992), South Africa (Nickles et al., 2026), Colombia (Vargas et al., 2019), Chile (Nouhra, Palfner, Kuhar, Pastor, & Smith, 2019) and Argentina (Giles, Salomón, Pildain, & Barroetaveña, 2025).

Significant genetic variation exists within *A. muscaria*, which is considered to represent a species complex (Geml, Tulloss, Laursen, Sazanova, & Taylor, 2008). Multiple phylogenetic analyses have recovered three major clades that correspond geographically to Eurasian, Eurasian subalpine and American groups, which all overlap in Alaska (Geml et al., 2006, 2008; Oda, Tanaka, & Tsuda, 2004). The recognised variation in *A. muscaria* has led to the naming of several varieties and one species, *A. persicina*, with further subdivision of *A. muscaria* considered likely (Ferrer-Gallego et al., 2026; Tulloss et al., 2015). The Eurasian clade corresponds to *A. muscaria* var. *muscaria* (Nickles et al., 2026) and specimens from across the introduced range including Colombia, Argentina, South Africa, New Zealand and Australia all group with this Eurasian clade (Geml et al., 2006; Giles et al., 2025; Nickles et al., 2026; Vargas et al., 2019). Therefore, it appears that the Eurasian clade represents the primary taxa involved in the global spread and invasion of *A. muscaria* (Nickles et al., 2026).

Currently there are two highly contiguous reference genomes available for *A. muscaria* (Table 1). Both of these genomes represent the American clade, and include the varieties *A. muscaria* var. *formosa* (Lu et al., 2026); and *A. muscaria* var. *guessowii* (Hess et al., 2014). Two recent studies have published a number of less contiguous genomes (Table 1) from the Eurasian clade (Nickles et al., 2026; Su et al., 2026). These are highly valuable and informative for a range of specific questions (see (Nickles et al., 2026)), but are not contiguous enough to be a general reference for the clade. Given the phylogenetic divergence between the American and Eurasian clades, a contiguous reference genome for the Eurasian clade would be valuable for facilitating genomic analysis of ecological and evolutionary questions in *A. muscaria* var. *muscaria* across its exotic and natural ranges. To fill this gap in genomic resources, we assemble a high quality reference genome for *A. muscaria* var. *muscaria* from a New Zealand sporocarp.

**Table 1.** Existing genome resources in *Amanita muscaria*.

| Name | Koide Hess et al. (2014) | JGI Lu et al. (2026) | Su et al. (2026) | Nickles et al. (2026) |  |
|---|---|---|---|---|---|
|  |  |  |  | Average | Range |
| Location | Pennsylvania, USA | Washington, USA | China | USA, South Africa, Europe, Aus |  |
| Seq. method | Illumina HiSeq 2000 | PACBIO_ SMRT (Sequel II) | Illumina NovaSeq 6000 | Illumina NovaSeq 6000 |  |
| Size (bases) | 40.7M | 110.3M | 26.8M | 30.2M | 26.8M-41.1M |
| # scaffolds | 1101 | 316 | 2922 | 8020 | 24852-21336 |
| N50 | 168k | 2171k | 20k | 15k | 5k-24k |
| BUSCO | 95.6% | 97.5% | 91.7% | 92.0% | 75%-94.6% |
| NCBI Reference | GCA_ 000827485.1 | GCA_ 054369615.1 | - | - |  |
| JGI Reference | Amamu1 | Amaapr1 | - | - |  |

## Materials & Methods

### Sample details

The reference specimen was collected from Flagstaff forest, in the southern South Island of New Zealand (latitude: −45.82, longitude: 170.45) as a sporocarp on March 12th 2023. This site is a commercial *Pinus radiata* plantation owned by City Forests which was planted in 1995. Tissue samples were taken from the sporocarp cap tissue, avoiding gills and the outside surface to avoid spores and external contamination. These were snap frozen with liquid Nitrogen, and stored at −20°C.

### DNA extraction and sequencing

The sample was sequenced twice; with short read Illumina sequencing and with long read Oxford Nanopore Technologies (ONT) sequencing. For the short reads, one frozen tissue sample was ground as close as possible into a fine powder, using liquid nitrogen to keep the sample frozen during the grinding process. DNA was extracted from the collected tissue using a DNeasy^®^ Plant Mini kit (QIAGEN). The protocol was followed, except that only 50 *µ*L of elution buffer was used to re-suspend the DNA at the elution step. The DNA concentration in the resultant solution was measured with a spectrophotometer (NanoDrop™, Thermo Fisher Scientific). Samples were then con-centrated using a speedvac until the estimated DNA concentration exceeded 10 ngL^−1^. Whole genome resequencing was carried out by Novogene Singapore. 22 *µ*L aliquots of extracted DNA were transferred to fresh tubes which were sealed with parafilm and sent to Novogene Singapore on ice. DNA was sequenced with NEBNext Lib Prep Kit and Illumina NovaSeq 6000 PE150.

For the long-read sequencing, high molecular weight (HMW) DNA was obtained using a CTAB extraction adapted from that of Sahu, Thangaraj, and Kathiresan (2012). The frozen tissue sample was divided into six portions, freeze-dried overnight, then macerated in liquid nitrogen with a pestle to form a smooth powder. 750 *µ*L of CTAB (cetyltrimethylammonium bromide) buffer was added to each tube and left to incubate for 1 hour at 65°C. 400 *µ*L of chloroform was added to each tube, the mixture put on a tilt plate for 1 minute to mix and then centrifuged for 5 minutes at 12000G before the aqueous phase was transferred to a new tube. The chloroform step was repeated, and the aqueous phases from three extractions were combined to form two samples. 3.3 mL of 100% ethanol and 165*µ*L of sodium acetate were added to each sample. No visible pallet was observed at this stage. The mixtures were incubated for 30 minutes at −20°C, then centrifuged for 15 minutes at 12000G. A small amount of precipitate was observed. The supernatant was poured off, 5 mL of 70% ethanol added to each sample, and the samples were centrifuged for 2.5 minutes at 12000G. The ethanol was poured off, the tubes centrifuged for a few seconds to collect any remaining ethanol in the bottom of the tube, and this ethanol was removed by pipette. The precipitate was left to air dry at room temperature. DNA was resuspended overnight in 100 *µ*L of a 1:10 dilution of TE buffer. Finally, RNAse A was added at a ratio of 1*µ*L RNAse A to 100 *µ*L of sample and incubated for 50 minutes at 37°C. Both samples were combined. The sample was purified with two rounds of DNA clean-up using AmPure^®^ XP beads (Beckman Coulter) at 0.8 volume.

The extracted HMW genomic DNA (gDNA) was quantified on a Qubit™ fluorimeter with dsDNA HS Assay Kit (Thermo Fisher Scientific), while the A260/A280, A260/A230 ratio was assessed on the NanoDrop™ 2000 spectrophotometer (Thermo Fisher Scientific) and the estimated size was visualised on a 0.8% agarose gel. The gDNA was prepared into sequencing libraries using the “Ligation sequencing DNA” protocol for the Ligation Sequencing Kit 114 (SQK-LSK114; ONT). Approximately 25-30 femtomole (fmol) of the library was loaded onto a FLO-MIN114 (R10.4.1) Flow Cell and sequenced for 72 h on the ONT GridION sequencer in the Otago Genomics Facilty. A total of two loads of library were sequenced, with one wash to recover pores in between loading. Basecalling was performed live with Guppy 7.1.4 (ONT) running on the MinKNOW software (23.07.12; ONT) at super-accurate basecalling mode (minimum Q-score threshold of 10).

### Genome assembly

The raw long reads were trimmed and filtered by quality. Reads with an average phred quality score less than 10 or strings of more than 240 consecutive A bases were removed using chopper 0.5.0 (Coster & Rademakers, 2023). Reads were trimmed using cutadapt (Martin, 2011) to remove sequences of bases at the start and/or end of each read with a quality score less than 10. The correction function of Canu 2.2 (Koren et al., 2017) was used to create an error-corrected subset of the trimmed and filtered reads with 40*×* genome coverage. Short reads from the same individual were trimmed and filtered with cutadapt (Martin, 2011), removing adapters, ends of reads with a phred score less than 25, and reads shorted than 50 bp.

Raw nanopore long reads were assembled using Flye 2.9.1 (Kolmogorov, Yuan, Lin, & Pevzner, 2019), with genome size 40M. This was polished with medaka 1.6.0 (Oxford Nanopore Technologies Ltd, 2018), with the model specified as r941 min sup g507, first using the raw nanopore reads and then using the trimmed, filtered and corrected nanopore reads. This was followed by polishing with the trimmed and filtered short reads using Pilon 1.24 with default settings (Walker et al., 2014). The resulting assembly was scaffolded with LongStitch 1.0.4 (Coombe et al., 2021) using the trimmed, filtered nanopore reads. Settings G=3.6e7, gap fill=True, c=2 and l=2 specified the expected genome size and the requirements for filling gaps. Two rounds of LongStitch were run; the first with the default pipeline of Tigmint (Jackman et al., 2018) and ntLink (Coombe, Warren, Wong, Nikolic, & Birol, 2023), and the second round with the ntLink-ARKS (Coombe et al., 2018) pipeline for extra scaffolding.

Contaminants were removed using fcs 0.5.0 (Astashyn et al., 2023). Duplicate contigs and alternate haplotigs were removed with purge dups 1.2.6 (Guan et al., 2020). Approximately two thirds of the contigs were purged in this step. A second round of polishing with Pilon 1.24 (Walker et al., 2014) using the trimmed, filtered short reads followed.

A second draft assembly was created using the raw short reads with MaSuRCA 4.1.0 (Zimin et al., 2013). The long read and short read assemblies were combined using quickmerge (Chakraborty, Baldwin-Brown, Long, & Emerson, 2016) to produce a hybrid genome assembly. The long read assembly was treated as the primary assembly and the short read assembly was treated as the reference, so the short read assembly provides mostly the sequences which bridge the gaps in the “primary” long read assembly. This produces a more contiguous assembly than the alternative configuration of primary and reference assembly, and makes the most of the high quality assembly and long read structural data which has gone into the long read assembly. Duplicate sequences were removed with purge haplotigs 1.1.2 (Roach, Schmidt, & Borneman, 2018) with low/mid/high parameters 5*/*25*/*60, using the short reads. Finally, the hybrid assembly was polished with Pilon 1.24 (Walker et al., 2014) using the trimmed, filtered short reads.

The mitogenome was assembled separately to maximise the chance of a complete assembly. An assembly graph was constructed using the trimmed, filtered short reads and SPAdes 3.15.4 (Prjibelski, Antipov, Meleshko, Lapidus, & Korobeynikov, 2020). This was fed into GetOrganelle 1.7.7.0 (Jin et al., 2020) and the fungal mitochon-

dia database was used to extract and assemble the mitogenome. This mitogenome was blasted 2.13 (Altschul, Gish, Miller, Myers, & Lipman, 1990) against the hybrid genome assembly. One scaffold of the hybrid assembly was identified as matching parts (approximately 10%) of the mitogenome and was replaced with the separately assembled mitogenome.

Repeats were masked with RepeatMasker 4.1.0 (Smit, Hubley, & Green, n.d.), specifying that the genome belongs to Agaricomycetes. The genome was annotated with AUGUSTUS 3.3.3 using the model trained on *Laccaria bicolor* (Hoff & Stanke, 2013; Stanke, Diekhans, Baertsch, & Haussler, 2008). Coverage of conserved genes was evaluated with BUSCO 5.1.3, using the fungi odb10 lineage (Manni, Berkeley, Seppey, Simão, & Zdobnov, 2021). Assembly statistics were calculated with gfastats 1.3.6 (Formenti et al., 2022).

### Phylogenetic placement

A phylogeny was constructed to confirm the taxonomic identity of the newly sequenced specimen. The DNA sequences of the ITS, LSU and *β*-tubulin regions from the samples used in Geml et al. (2006) were downloaded from NCBI. The ITS, LSU and *β*-tubulin regions of the newly assembled genome, the Koide BX008 assembly GCA 001691765.1 and the *A. muscaria* var. *formosa* assembly from JGI were identified with BLAST (Altschul et al., 1990) and their sequences were retrieved. An out-group was included for the ITS and *β*-tubulin regions; a sample of *A. pantherina* from Oda et al. (2004) (NCBI IDs: AB080978 and AB095867). For each gene, all samples were aligned with MAFFT (Katoh, 2013). The three gene sequences for each sample were concatenated, with missing data inserted where appropriate, and a maximum likelihood phylogeny was constructed with IQ-TREE 2.2.2.2 (Minh et al., 2020), using ModelFinder (Kalyaanamoorthy, Minh, Wong, von Haeseler, & Jermiin, 2017) to select the best fit substitution model. A consensus tree was constructed using 1000 bootstrap replicates (Hoang, Chernomor, von Haeseler, Minh, & Vinh, 2018).

## Results & Discussion

The Oxford Nanopore sequencing basecalling generated 9.03 Gb of sequence data, representing approximately 240*×* read depth for the entire genome. This data comprised 2.1 million reads with an N50 of 6.92 kb (i.e. 50% of the bases are in reads of 6.92 kb or longer). The mean length of the Nanopore reads is 4, 322.3 bp and the overall quality was good relative to the approach with a mean phred score of 16.4, which relates to a per-base error probability of about 2.3%. Illumina sequencing resulted in 759369 unique forward and 821084 unique reverse reads. After trimming, the GC content is 47% and no sequences were of poor quality.

A haploid genome assembly was produced, comprising 240 nuclear contigs and one circular mitogenome. The final assembly is composed of 241 scaffolds with a total length of 39, 041, 218 base pairs. It has an N50 of 371, 573 (N50 is the size of the scaffold where at least half of the assembly is in scaffolds of that length or longer [larger = better]) and an L50 of 28 (L50 is smallest number of scaffolds that contain at least half of the bases in the assembly [smaller = better]). These statistics measure the most contiguous half of the genome; L50 is smallest number of scaffolds which can be chosen to contain at least half of the bases in the assembly and N50 is the size of the smallest scaffold such that at least half of the assembly is contained in scaffolds of that length or longer. Further details are provided in Table 2. The assembly has a mean long read depth of 192*×*, and 86% of assembled bases are covered by at least 100 long reads.

**Table 2.** Summary statistics of the hybrid genome assembly, including the mitogenome.

| Statistic | Value |
|---|---|
| # scaffolds: | 241 |
| Total scaffold length: | 39,041,218 |
| Average scaffold length: | 161,996.76 |
| Scaffold N50: | 371 573 |
| Scaffold auN: | 539 269.92 |
| Scaffold L50: | 28 |
| Largest scaffold: | 1,953,853 |
| Smallest scaffold: | 1063 |
| # contigs: | 319 |
| Total contig length: | 38,465,120 |
| Average contig length: | 120,580.31 |
| Contig N50: | 256,641 |
| Contig auN: | 363,696.53 |
| Contig L50: | 40 |
| Largest contig: | 1,271,339 |
| Smallest contig: | 704 |
| GC content %: | 47.58 |

Genome annotation and gene prediction (using AUGUSTUS) estimated that the genome contains 11752 genes.

Coverage of conserved fungal genes was evaluated with BUSCO. Of the 758 BUSCO groups common to fungi, 95.1% were found in this genome; 93.1% as a single copy (Table 3). Repeat masking masked 217, 948 base pairs, comprising 0.56% of the genome. Most of the masked regions were simple repeats, and the remainder were low complexity sequences.

**Table 3.** BUSCO results for genes present in the *Amanita muscaria* genome assembly (with the mitogenome included) from dataset fungi odb10.

| Number | Proportion | Type |
|---|---|---|
| 721 | 95.1% | Complete BUSCOs |
| 706 | 93.1% | Complete and single-copy BUSCOs |
| 15 | 2.0% | Complete and duplicated BUSCOs |
| 14 | 1.8% | Fragmented BUSCOs |
| 23 | 3.1% | Missing BUSCOs |
| 758 | 100% | Total BUSCO groups searched |

Phylogenetic analysis using the ITS, LSU and *β*-tubulin genes showed that, as we predicted, the new NZ genome groups with the *A. muscaria* Clade II described by (Geml et al., 2006) (Figure 1). In particular, our genome groups with a sample from

**Figure 1.**
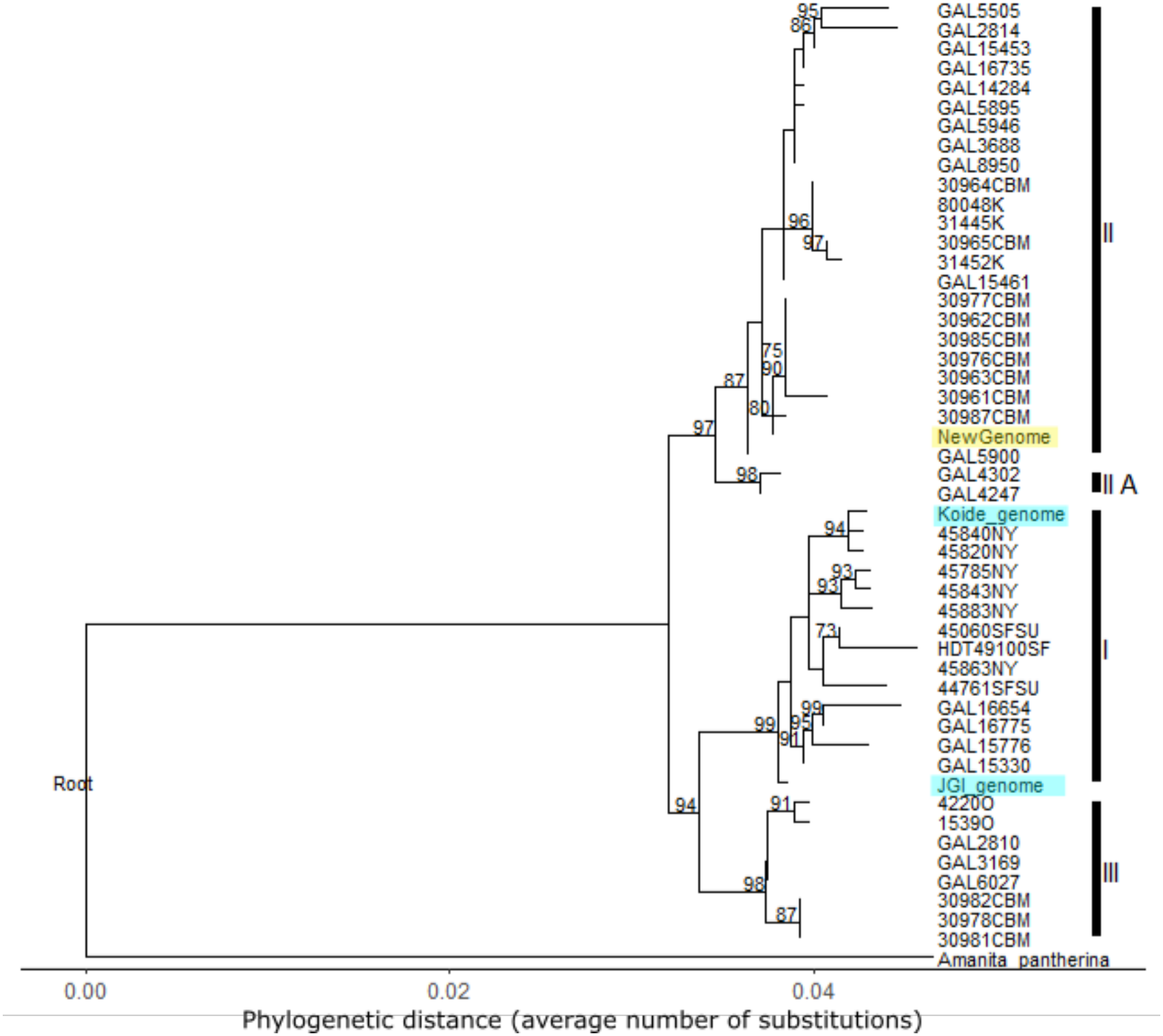
Synteny map between the new NZ genome assembly described here (x-axis) and the Koide BX008 assembly (y-axis). Gridlines represent scaffolds.

Queenstown, New Zealand, and with samples from Japan (Geml et al., 2006). The Koide BX008 genome and the *A. muscaria* var. *formosa* genome (labelled JGI) both sit within the North American Clade (Clade I of (Geml et al., 2006)). These findings are consistent with previous work showing that *A. muscaria* in its exotic range is to date exclusively represented by the Eurasian Clade, and the taxon *A. muscaria* var *muscaria*. This confirms the utility of our genome as the most contiguous genomic resource for this globally significant taxa Our new NZ *A. muscaria* genome assembly compares well to the two existing contiguous reference genomes from North American Clade I (Table 4). The new NZ assembly is more complete and contiguous than the Koide BX008 assembly, which has N50 and L50 values that are indicative of a more fragmented assembly (scaffold N50 of 168 kb and L50 of 54 for Koide BX008 vs N50 of 372 kb and L50 of 28 for the new NZ assembly), over ten times the number of contigs, and four times as many scaffolds. The JGI var. *formosa* assembly is more comparable in terms of continuity statistics but is just under three times the size of our assembly, and much larger than the estimated genome size for *A. muscaria*. This expanded size may be indicative of assembly problems such as failure to collapse alternative haplotypes, or contamination (Guiglielmoni, Houtain, Derzelle, Van Doninck, & Flot, 2021). We investigated this size difference further using synteny mapping to compare our new NZ genome to both the JGI var. *formosa* and the Koide BX008 assemblies. We found high levels of synteny between our genome and the Koide BX008 assembly but a much noisier relationship with the JGI var. *formosa* genome (Supp. fig. 1, 2). A restricted synteny map looking at the largest 28 scaffolds shows that most of our scaffolds map to two locations on the JGI var. *formosa* genome (Supp. fig. 3) indicating that failure to collapse haplotypes is likely to be partly driving the size of the JGI var. *formosa* assembly. Uncollapsed haplotypes are a well-known issue in assemblies of non-model organisms and can lead to structurally incorrect assemblies (Guiglielmoni et al., 2021). Structural inaccuracies would be consistent with the very noisy synteny results between our genome and the JGI var. *formosa* assembly.

**Table 4.**
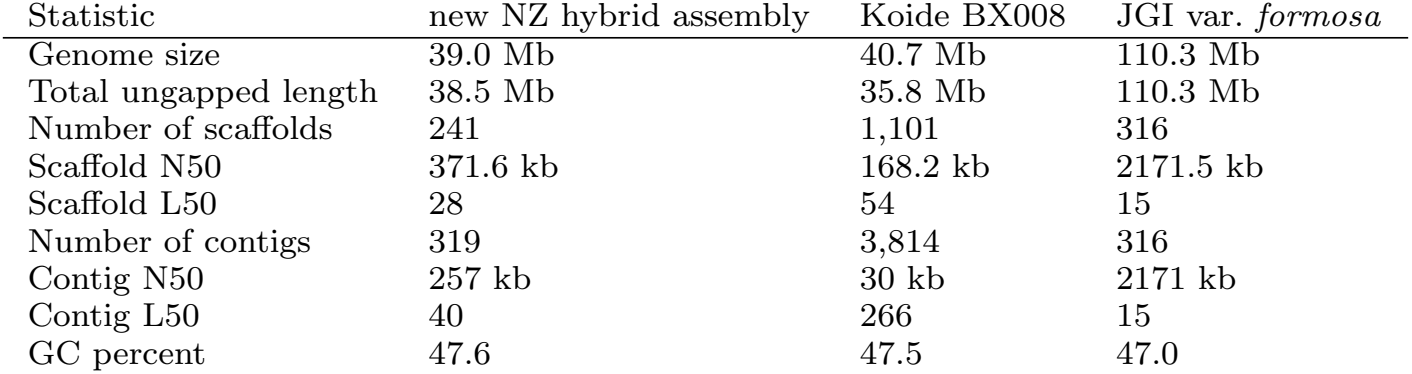
Comparison of summary statistics for genome assemblies of *Amanita muscaria*. The new NZ assembly is the hybrid assembly with the separately assembled mitogenome, as described above. The Koide BX008 assembly is hosted on NCBI with GenBank reference GCA 000827485.1 and the var. *formosa* assembly is available on JGI as *Amanita muscaria* var. *formosa* 2016PMI152 v1.0. Statistic new NZ hybrid assembly Koide BX008 JGI var. *formosa*

**Figure 2.**
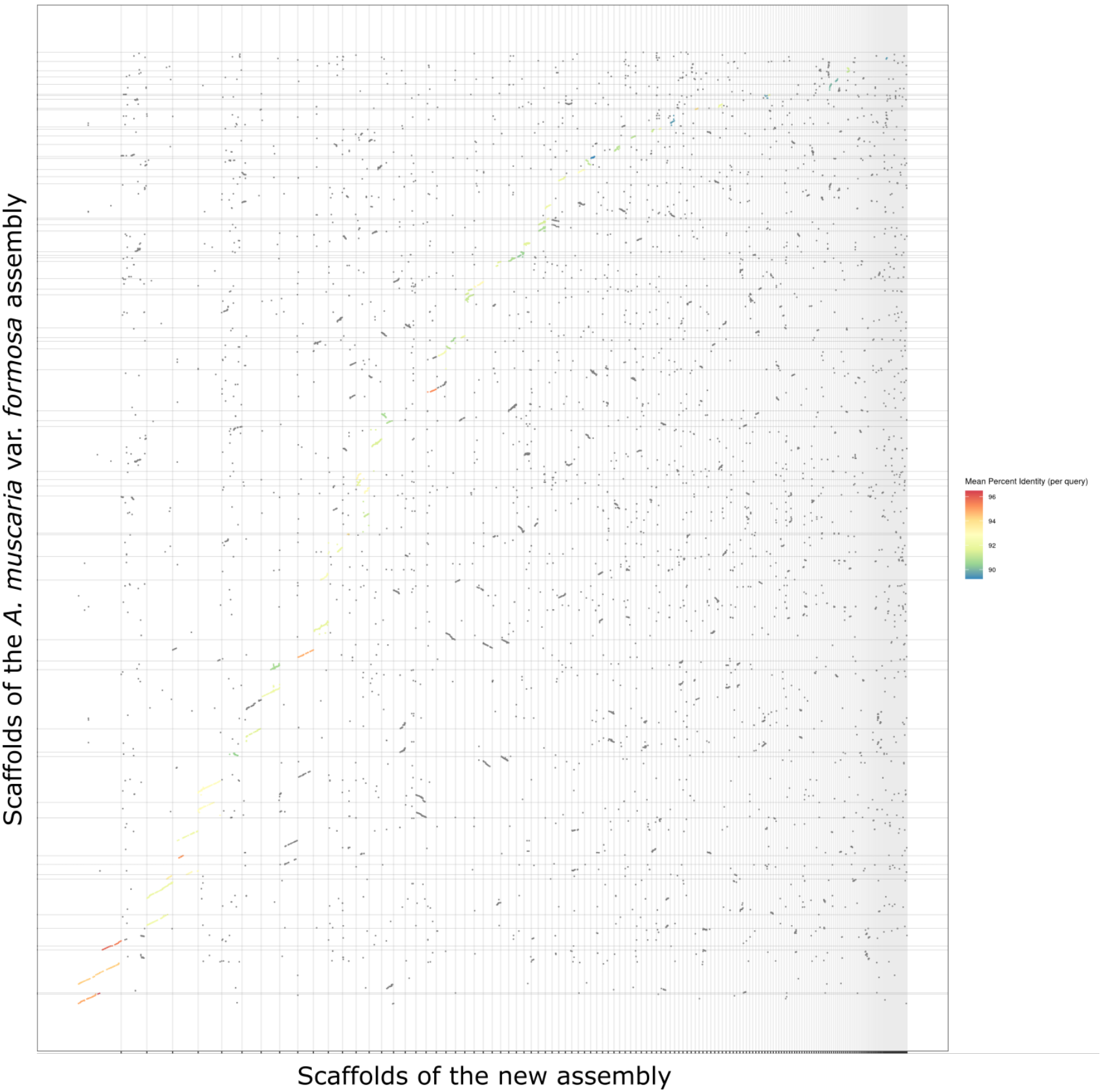
Synteny map between the new NZ genome assembly described here (x-axis) and the JGI var. *formosa* assembly (y-axis). Gridlines represent scaffolds.

**Figure 3.**
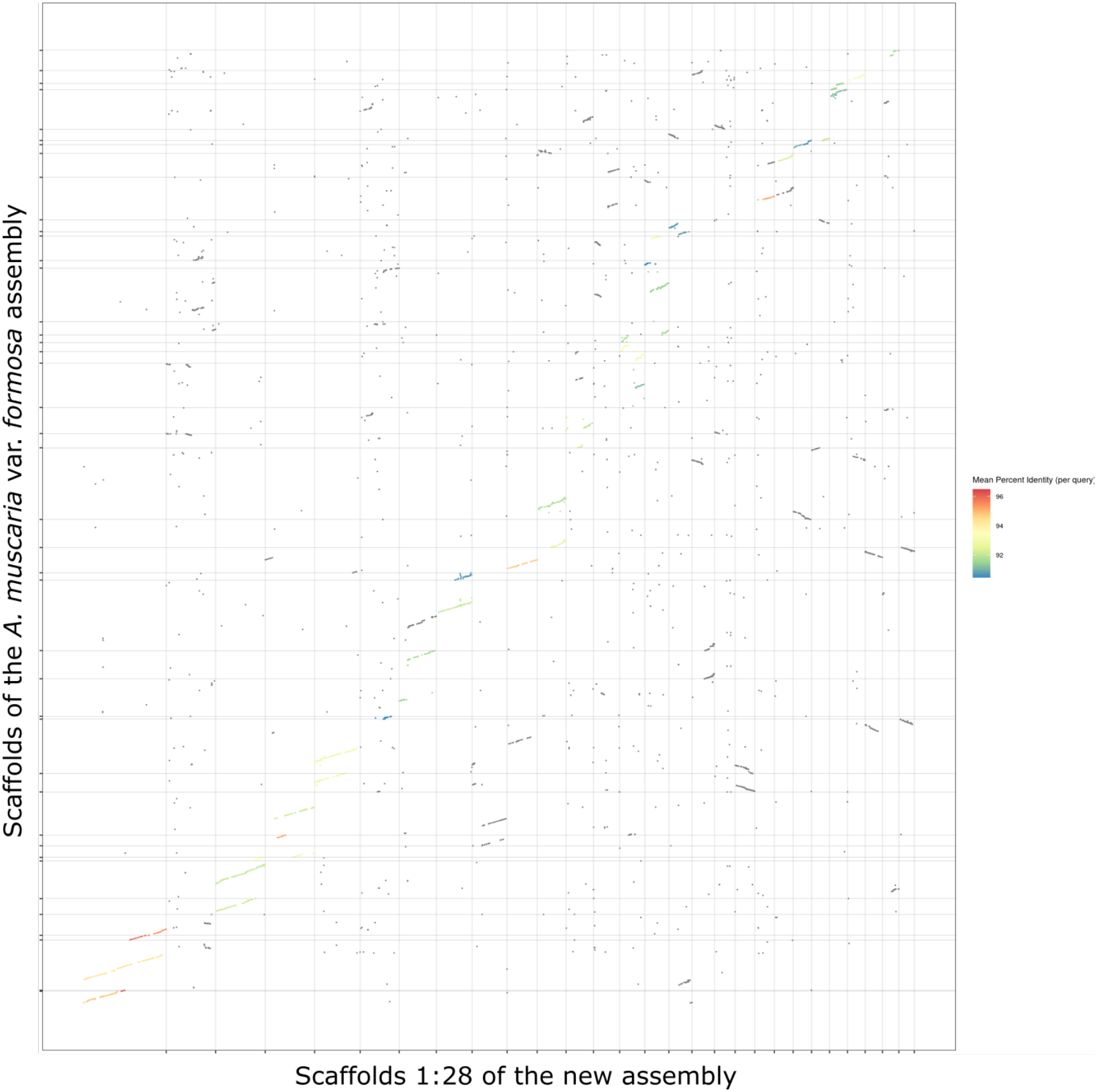
Restricted synteny map between the new NZ genome assembly described here (x-axis) and the JGI var. *formosa* assembly (y-axis), restricted to the largest 28 scaffolds of the new NZ assembly. Gridlines represent scaffolds.

The new NZ genome presented here represents the highest quality resource available for *A. muscaria* in the broad senses, and is also comparable to other *Amanita* assemblies (Hess et al., 2014; Pulman, Childs, Sgambelluri, & Walton, 2016; Zhou et al., 2021). Our new NZ genome will be a useful resources for annotating genes, especially hard to annotate regions such as mating type loci that are under balancing selection (Nieuwenhuis et al., 2013). It will also provide improved confidence in functional and comparative genomic analyses where the higher continuity will help better resolve links between genes and regulatory regions (Whibley, Kelley, & Narum, 2021), as well as improve insights into structural evolution among genomes (Manley et al., 2023). The new NZ *A. muscaria* var *muscaria* genome assembly has been submitted to the NCBI genome database with accession number JCGJAC000000000.

## Note

A new genome of *A. muscaria* (assemblies gfAmaMusc1.hap1.1 and gfAma-Musc1.hap2.1, GenBank accession GCA 988426935.1) has recently been released on NCBI by the Wellcome Sanger Institute. This new release is a chromosome-level assembly (12 chromosomes and one mitogenome, 50.3 Mb, scaffold N50 4.1 Mb) and is haplotype resolved. Extending the phylogenetic analysis described in this paper to include the new genome shows that it should belong to the European Clade II. This new release will supercede the genome described in this paper as a high-quality genomic reference.

## Funding

This work was supported by a Otago Doctoral Scholarship (to LT). The research was partly funded by Ministry of Business, Innovation, and Employment of New Zealand through a Data Science Programmes Grant (UOAX1932).

The authors wish to acknowledge use of the eResearch Infrastructure Platform hosted by the Crown company, Research and Education Advanced Network New Zealand (REANNZ) Ltd., and funded by the Ministry of Business, Innovation & Employment. URL: https://www.reannz.co.nz

## Data availability

The data in this work has been submitted to the NCBI database under the BioProject PRJNA1533062. The sequence reads are on the NCBI SRA database under accession numbers SRR40790224 and SRR41509576. The genome has accession number JCGJAC000000000.

## Additional figures

**Figure 1.**
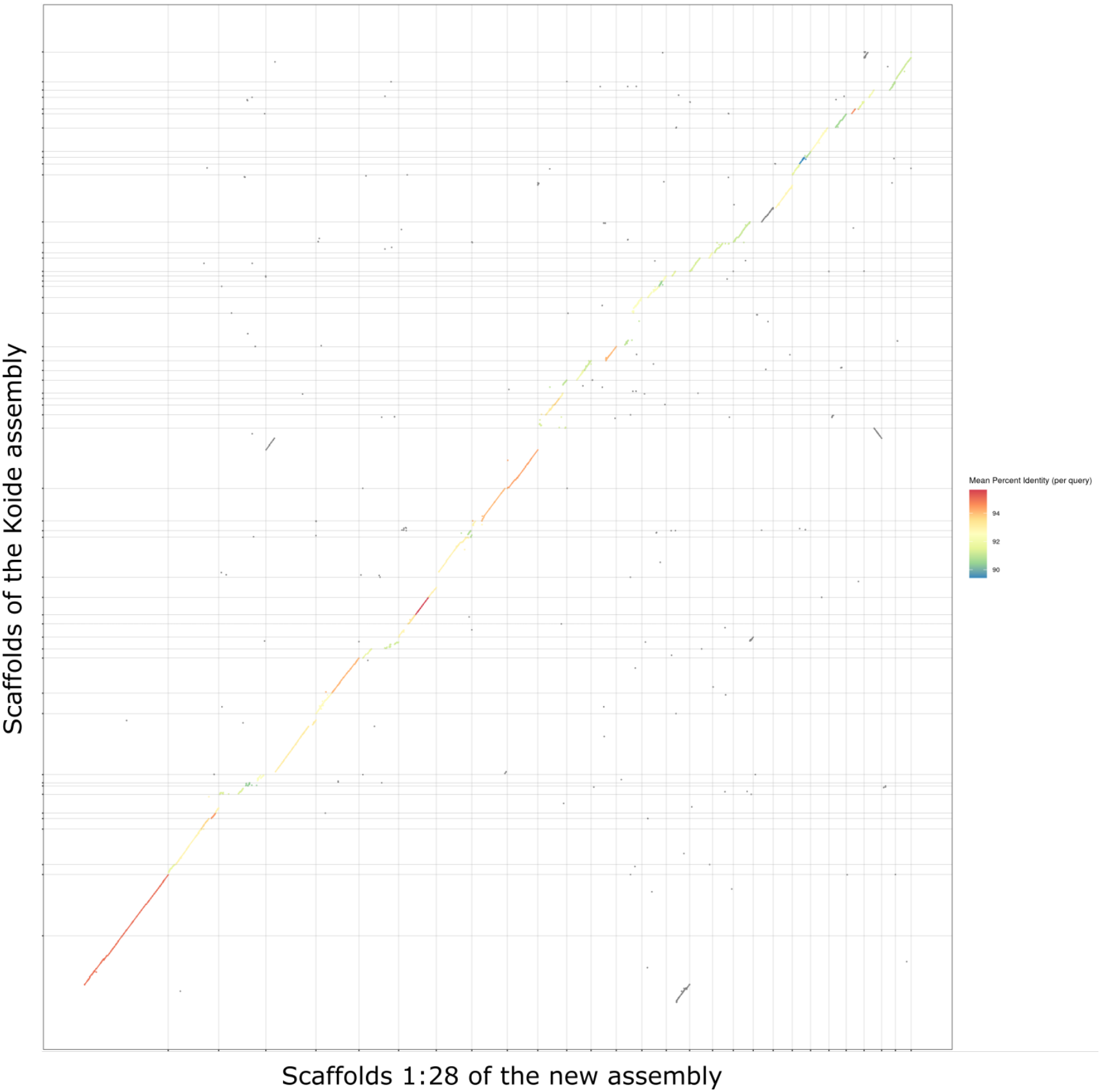
Consensus tree based on the ITS, LSU and *β*-tubulin genes of *Amanita muscaria*, and generated from 1000 bootstrap replicates of maximum likelihood trees. Bootstrap values are shown where greater than 70%. Clade labels on the right reflect the clades identified by Geml et al. (2006). The new NZ genome of this paper (yellow) and the two strains with genome assemblies (blue) are highlighted.

## Notes

### Competing Interest Statement

The authors have declared no competing interest.

